# Differences in the temporal processing between identification and categorization of durations: a behavioral and ERP study

**DOI:** 10.1101/315887

**Authors:** Dorian Bannier, J. Wearden, Christophe C. Le Dantec, Mohamed Rebaï

## Abstract

This study examined how different forms of decision-making modulate time perception. Participants performed temporal bisection and generalization tasks, requiring them to either categorize a stimulus duration as more similar to short or long standards (bisection), or identify whether or not a duration was the same as a previously-presented standard (generalization). They responded faster in the bisection task than in the generalization one for long durations. This behavioral effect was accompanied by modulation of event-related potentials (ERPs). More specifically, between 500 ms and 600 ms after stimulus offset, a late positive component (LPC), appearing in the centro-parietal region, showed lower amplitude in the bisection task than in the generalization one, for long durations, mirroring the behavioral result. Before (200-500 ms) and after (600-800 ms) this window, the amplitude of the LPC was globally larger in the generalization paradigm, independently of the presented duration. Finally, the latency of the LPC’s peak was earlier for long durations than for the short ones, indicating that the decision about the former stimuli was made earlier than for the latter ones. Taken together, these results indicate that the categorization of durations engages fewer cognitive resources than their identification.

## 1. Introduction

Pacemaker-accumulator models of time perception propose that durations are encoded by an internal clock before being stored in memory and evaluated by decision-making processes (Gibbon, Church, & Meck, 1984; Wearden, 2005). This cognitive structure can be appropriately understood using the common situation encountered while waiting at a traffic light. When we stop at the red light, we first encode its duration using our internal clock. The representation resulting from this encoding is stored in working memory in order to be compared, in real-time, with a reference duration of similar experiences stored in long-term memory. Then, decision-making processes allow us to determine if the current red light duration is abnormally long or to anticipate the green light.

Many experimental tasks have been designed to probe time perception (Grondin, 2010). It is possible to ask participants to compare pairs of consecutive stimuli, and to report which one is the shorter (e.g.Le Dantec et al., 2007) or whether their durations are the same (Wearden & Bray, 2001). Another way to test time perception consists of comparing a presented duration with a previously learned one. This method includes two paradigms widely used in time perception research: the generalization and the bisection tasks. In a generalization task, participants have to determine if a presented duration is the same or not as a standard duration learned at the beginning of the experiment (Kononowicz & van Rijn, 2014; Paul et al., 2011; Wearden, 1992). Hence, this protocol requires identification of the standard interval. In a bisection task, subjects have to determine if a presented duration is closer of one of two previously learned short or a long standard durations (Ng, Tobin, & Penney, 2011; Wearden, 1991; Wiener & Thompson, 2015). Therefore, participants categorize intervals in this paradigm.

Modelling temporal processing in generalization and bisection tasks has sometimes emphasized potential similarities in the decision-making involved in the two tasks, but also sometimes suggested differences. Wearden (1992) proposed that, in the generalization paradigm, a comparison is made between the presented duration and the standard. If the difference is sufficiently large, and exceeds a threshold, participants report that the presented duration is different from the standard one. Otherwise, they respond that it is the standard.

Some accounts of bisection (e.g.Wearden, 1991) propose that this task involves two comparisons, one between the current duration and the short standard, and the other between the current duration and the long standard, with the obvious suggestion that bisection is more cognitively complex than generalization in terms of the comparison and decision processes required. On the other hand, Wearden (2004) proposed that, in the bisection paradigm, a duration criterion, which corresponds to the mean of all the durations presented in the experimental set, may be formed (see also Wearden & Ferrara, 1996). If a presented duration is sufficiently lower or higher than the criterion duration, a “short” or “long” response is made. Hence, in the two cases, it may be that responses are based on a comparison between a presented duration and a reference value that is supposed to be the center of the distribution of the durations presented to the participants, thus rendering the tasks more similar in terms of cognitive complexity.

However, some more recent work has also suggested that temporal generalization and bisection are not based on similar processes, for different reasons. Ogden, McKenzie-Phelan, Montgomery, Fisk, and Wearden (2018) examined performance on two types of generalization and bisection tasks. One type was “normal”, for example, standard durations were presented at the start of the study, and refreshed from time to time. The other type was “episodic”: in temporal generalization two stimulus durations were presented on each trial, and the task was to judge whether or not they had the same duration. In bisection, three stimuli were presented, a short and long value, and a stimulus to be judged. The current stimulus had to be classified in terms of its similarity to the short and long values presented on the trial, but these were different on each trial (see Wearden & Bray, 2001). The basic idea of the experiment was to compare performance on the “normal” version of the tasks, which supposedly involved reference memory of one or two standards, and the “episodic” versions which did not, as stimulus durations changed on each trial. For present purposes, the important result was that performance on the “normal” and “episodic” temporal generalization tasks differed, whereas this was not true for the bisection task: the versions supposedly with and without reference memory of standards produced the same behavior. Ogden et al. (2018) did not present a formal model of performance on bisection, but a potential conclusion from their study is that bisection is based on some “local” criterion, which may vary from trial to trial. See also Droit-Volet and Rattat (2007)for a similar idea.

In general, event-related potential (ERP) studies have revealed differences between these two tasks, perhaps reflecting the qualitatively different responses made during the categorization and the identification of durations, and suggesting that they are not performed in the same way. These studies mostly relied on the investigation of the contingent negative variation (CNV), a fronto-central component extensively studied in time perception (Kononowicz, van Rijn, & Meck, 2016; Ng & Penney, 2014). The CNV is a slow negative wave developing across time between the beginning and the end of a presented duration. Although its direct link to temporal processing had been debated, the fact that the CNV develops across time means that this activity is somehow indirectly guided by temporal information (Kononowicz &van Rijn, 2011; van Rijn, Kononowicz, Meck, Ng, & Penney, 2011). In this context, it has been shown that in bisection task and for long durations, the amplitude of the CNV develops until the short standard duration has elapsed, then the negativity is constant until about the geometric mean of the distribution of durations presented (Ng et al., 2011). In the generalization task, the development of the CNV appears constant across the entire duration presentation. Moreover, the generalization task appears to differently mobilize the two cerebral hemispheres. For long durations, the amplitude of the CNV increased until the equivalent of the standard duration had elapsed in the left hemisphere, while it continued to increase until the end of the presented duration in the right hemisphere (Pfeuty, Ragot, & Pouthas, 2003). Taken together, these results indicate differences in the cerebral dynamics supporting temporal processing during the categorization and the identification of durations.

Although differences can be inferred between the bisection and generalization tasks, no direct contrast has ever been made in order to compare them. Consequently, the aim of this study was to directly investigate how decision-making differs between the categorization and the identification of durations, where the stimulus durations involved were the same in the two tasks. Participants performed a bisection task where they had to categorize stimulus durations as short or long, and a generalization one, where they had to report whether or not presented durations were the standard. These protocols allowed us to contrast two different hypotheses. Firstly, if the bisection task relies on the comparison between the presented duration and the two standards (short and long), then the cognitive demands should be higher in this paradigm than in the generalization one that requires only a comparison between the presented duration and one standard. Secondly, if the bisection task does not require any long-term memory storage of standard durations, as previously suggested (Droit-Volet & Rattat, 2007; Ogden et al., 2018), then the generalization task should require more cognitive resources.

To test these two hypotheses, we used both a behavioral and an event-related potentials approach. As responses are different between the two paradigms, we investigated behavioral differences based on reaction times measured from the offset of the durations, in order to prevent the effect of physical duration of stimuli on temporal processing. Regarding the event-related potentials, the analyses focused on centro-parietal signals evoked after the offset of the duration. This region and this time window have previously been associated with attentional and working memory processes implicated in temporal processing (Lindbergh & Kieffaber, 2013), and are expected to be modulated by the differential cognitive demands of the generalization and bisection tasks, if these are different. If the bisection task engages fewer resources than the generalization one, reaction times should be shorter and the amplitude of the centro-parietal evoked potentials should be smaller in bisection than generalization. In contrast, if the bisection task engages more resources, the opposite pattern should be observed.

## 2. Methods

### 2.1. Participants

Twenty-one healthy participants with normal or corrected vision were recruited. All of the participants provided informed consent, and testing was conducted according to the declaration of Helsinki (BMJ 1991 ; 302 : 1194). They were all right handed, according the Oldfield’s criteria (Oldfield, 1971). Three participants were excluded from the ERP analysis because they presented less than 20 signals by condition. The final sample was composed of eighteen participants (including eight men; mean age: 21.06 ± 0.94).

### 2.2. Stimuli

The visual stimuli were composed of a white spot (0.5° of visual angle) presented in the center of a computer screen with a black background. The stimulus was displayed on a 15-inch screen with a resolution of 1280 × 1080 pixels and a refresh rate of 60 Hz.

### 2.3. Procedure

The participants performed a generalization and a bisection temporal task implemented with E-Prime^®^ software (Schneider, Eschman, & Zuccolotto, 2002). In the generalization task, they were instructed to indicate whether or not a presented duration was a standard duration (800 ms) previously learned. In the bisection task, the participants had to judge if a presented duration was closer to a short (200 ms) or to a long (1400 ms) standard duration, both previously learned. The task order was counterbalanced across participants. In both paradigms, participants were exposed to exactly the same set of stimulus durations. In each task, the durations used were: 200 ms, 400 ms, 600 ms, 800 ms, 1000 ms, 1200 ms, 1400 ms.

Each condition began with a training phase. In the generalization task, participants were exposed to 5 consecutive presentations of the standard duration (800 ms). They were told that they had only to memorize it, without responding. In the bisection task, 5 consecutive presentations of the short standard duration (200 ms) preceded 5 other consecutive presentations of the long standard duration (1400 ms). As in the generalization task, participants had to memorize them without responding.

The experimental phase followed this training phase in the generalization and in the bisection tasks. The experimental phase consisted of 27 blocks, each comprising a reminder of the standard(s) and the presentation of 21 comparison durations. The reminder was three consecutive presentations of the standard duration (800 ms) in the generalization paradigm, and three consecutive presentations of each of the standard durations (200 ms and 1400 ms) in the bisection one. Then, each of the seven comparisons durations (200 ms, 400 ms, 600 ms, 800 ms, 1000 ms, 1200 ms, 1400 ms) was presented three times in a random order. Between each trial, an empty screen followed for a random interval between 750 ms and 1250 ms. Hence, each task included 567 stimulus presentations, including 81 presentations of each of the durations and lasting approximately 30 minutes.

A practice session preceded the experimental phase just described, in order to familiarize participants with the task. This was a block that was exactly the same as the ones administered in the experimental session.

### 2.4. Behavioral analysis method

In the generalization task, a gradient was represented plotting the proportion of « yes, it is the standard » responses as a function of the presented duration. In order to test the asymmetry of the gradient, i.e. the fact that at a particular temporal distance from the standard duration, a duration longer than the standard one is more confused with it than a shorter one, « concentric » paired comparison was made using Wilcoxon tests (as in Ferrara, Lejeune, & Wearden, 1997). This procedure involved the comparison of the proportion of « yes » responses between 600 ms and 1000 ms, 400 ms and 1200 ms, 200 ms and 1400 ms.

In the bisection task, a logistic regression was used to fit a psychometric function, based on the proportion of « long » responses as a function of the presented durations. From this function, the bisection point, the duration for which the participants responded « long » in 50 % of trials, was calculated.

The behavioral comparison between the generalization and the bisection tasks was based on a reaction time (RTs) analysis. In order to eliminate the effect of physical duration on subjective temporal processing, the reaction times were measured from stimulus offset. RTs exceeding two standard durations were removed. A repeated-measures ANOVA was used to test the effect of duration (200, 400, 600, 800, 1000, 1200, and 1400 ms), task (bisection, generalization) and the interaction between these factors on the reaction times. Post-hoc comparisons involved Holm-Bonferroni correction for multiple comparisons and the Cohen’s ‘d’ was used to report a measure of the effect size. Between 0 and 0.2, the effect is considered negligible, small between 0.2 and 0.5, medium between 0.5 and 0.8, and large when higher (Cohen, 1992).

### 2.5. EEG acquisition and analysis methods

Continuous EEG data were acquired from 32 scalp electrodes (Waveguard, ANT, Enschede, The Netherlands), digitized with a sampling rate set at 512 Hz and with impendances kept below 10 KΩ. The recording used the following electrode set : Fp1, Fpz, Fp2, F7, F3, Fz, F4, F8, FC5, AFz, FC2, FC1, FC6, T6, C3, Cz, C4, T8, M1, CP5, CP1, CP2, CP6, M2, P7, P3, Pz, P4, P8, POz, O1, Oz and O2, with AFz being the reference electrode. Data were processed using ASA software. It was re-referenced offline using an average reference and filtered using high-pass and low pass filters (band-pass: 0.1-30 Hz). Trials were rejected if the absolute value of the signal amplitude exceeded ± 75 μV, on at least one of the electrodes.

Averaged ERP waveforms were computed for each of the durations in both paradigms, averaging between -100 ms before and 1000 ms after stimulus offset. Epochs were then baseline-corrected relative to the 100 ms interval prior to stimulus offset. Measures of mean amplitude were made on a centroparietal region of interest (ROI) including Pz, CP1, CP2, P3 and P4 electrodes. This ROI was chosen because after visual inspection, this region showed the highest amplitudes on the scalp. Moreover, this ROI was as close as possible to previous ones used in studies investigating temporal processing after the end of stimulus presentations (Lindbergh & Kieffaber, 2013; Macar & Vidal, 2003). As the ERPs evoked by the offset of stimuli resulted in long positive activities, we measured the mean amplitude in six consecutive 100 ms temporal windows from 200 ms to 800 ms post-offset.

A repeated-measures ANOVA was then performed in order to test the effect of Duration (200 ms, 400 ms, 600 ms, 800 ms, 1000 ms, 1200 ms, 1400 ms), Task (Bisection, Generalization) and the interaction between both factors on the mean amplitude in each of the six temporal windows. As for the behavioral analysis, post-hoc analysis used planned-comparisons and Holm-Bonferroni correction for multiple comparisons. Cohen’s ‘d’ was reported as a measure of effect size.

## 3. Results

### 3.1. Behavior

#### 3.1.1. Gradient in the generalization task

Figure 1 shows the temporal generalization gradient, in the form of the probability of a « Yes, it is the standard » response as a function of the presented durations. It appears that the curve peaked at the 800 ms standard duration but also to the 1000 ms comparison duration. In order to explore the asymmetry of the gradient traditionally demonstrated in human subjects, « concentric » pair comparisons (i.e. contrasting stimuli located at the same number of steps below and above the standard duration) were performed. Wilcoxon tests revealed that all three « concentric » pairs of stimuli produced significant effects. The 1000 ms, 1200 ms and 1400 ms durations gave rise to higher proportion of « yes » responses than 600 ms, 400 ms, and 200 ms, respectively (all p < 0.001).

**Figure 1:**
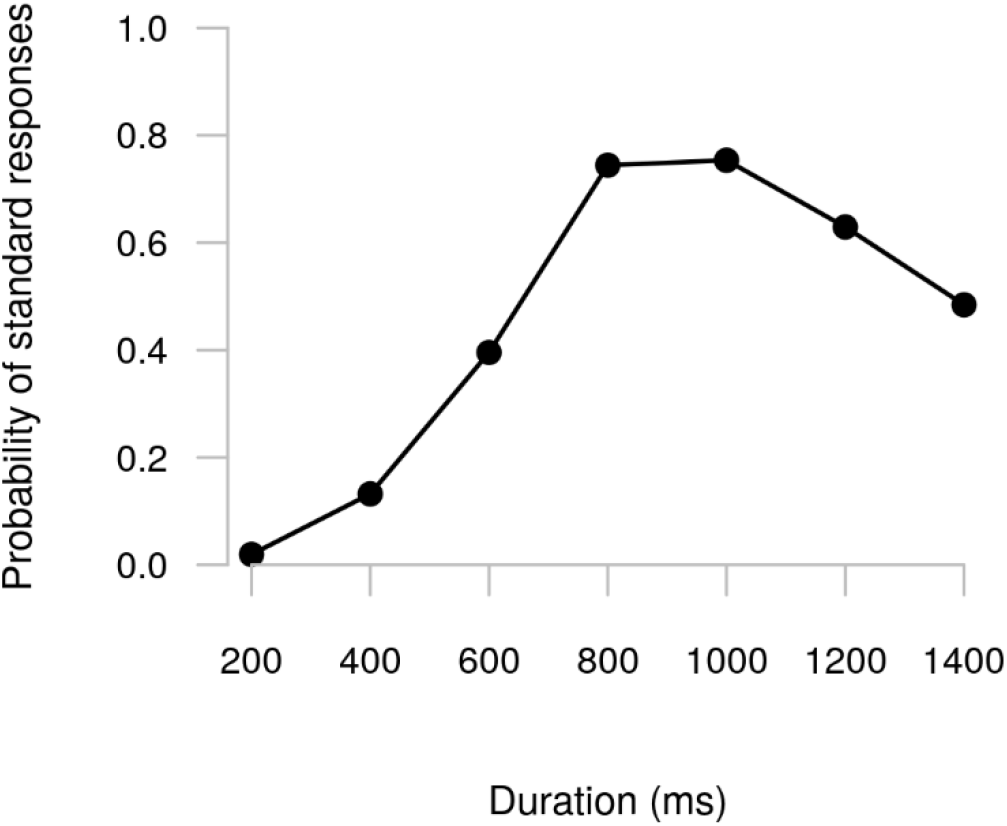
Probability of « standard » responses as a function of the presented duration in the generalisation task.

#### 3.1.2. Psychometric function in the bisection task

The figure 2 illustrates the psychometric function calculated by a logistic regression based on the probability of a « long » response in the bisection task. The regression analysis found the bisection point to be 722 ms (i.e. the duration at which participants responded « long » 50 % of trials).

**Figure 2:**
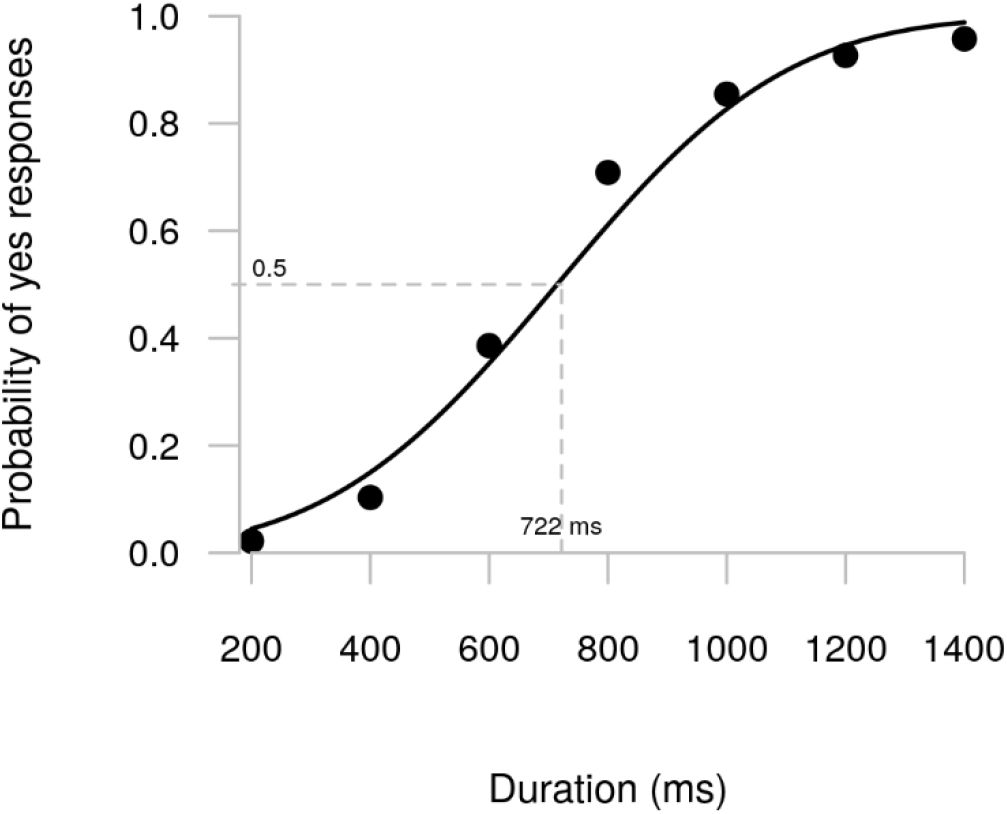
Probability of « long » responses as a function of the presented duration in the bisection task.

#### 3.1.3. Reaction times from offset

As participants had to respond to all durations in both paradigms, it was possible to explore their reaction times in both the generalization and bisection tasks. As mentioned in the method section, reaction times were measured from the offset of the stimuli, in order to control for the effect of physical duration on subjective time processing.

An ANOVA was performed using duration (200, 400, 600, 800, 1000, 1200 and 1400 ms) and task (generalization and bisection) as within-subject factors. This revealed significant main effects of duration (F(1,17) = 60.14 ; p < 0.001 ; ηp^2^ = 0.78) and of task (F(1,17) = 6.89 ; p = 0.018 ; ηp^2^ = 0.29), as well as a significant interaction between the two factors (F(1,17) = 6.91 ; p = 0.018 ; ηp^2^ = 0.29). Planned post-hoc comparison indicated that, for long durations, i.e. 1000 ms (t = 2.9 ; d = 0.59 ; p = 0.062), 1200 ms (t = 3.62 ; d = 0.77 ; p = 0.013) and 1400 ms (t = 2.79 ; d = 0.64 ; p = 0.062), RTs were shorter in the bisection task than in the generalization one (Figure 3). Albeit the effects were only tendencies for 1000 ms and 1400 ms, the Cohen’s d indicates that the effect of task for these durations was of medium size, as the effect size was between 0.5 and 0.8. Surprisingly, the same effect was obtained for 400 ms (t = 4.09 ; d = 0.5 ; p = 0.005).

**Figure 3:**
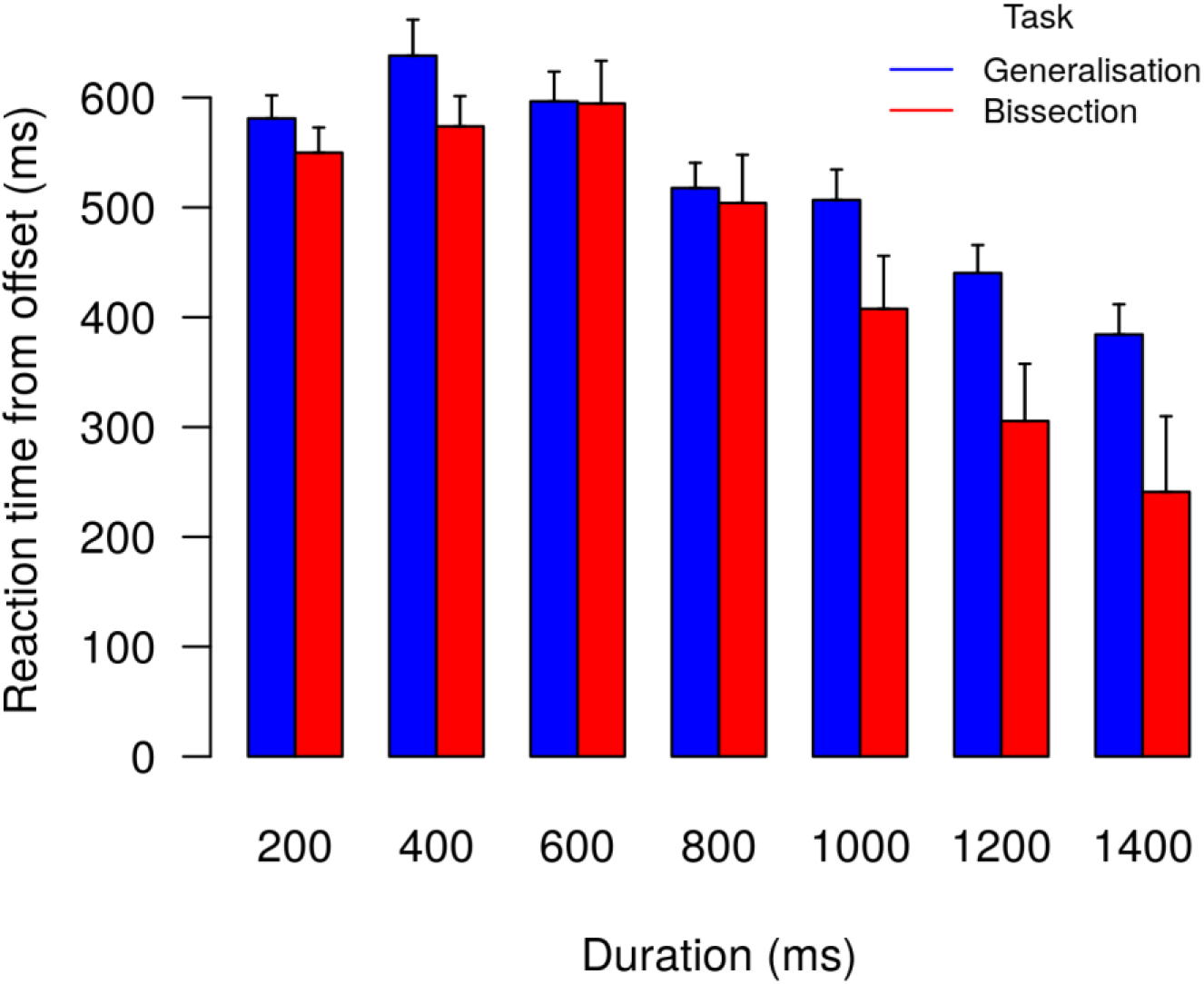
Probability of « long » responses as a function of the presented duration in the bisection task.

### 3.2. ERP’s

A large deflection was found occurring after each duration offset on centroparietal electrodes (Figure 4), consistent with previous findings (Lindbergh & Kieffaber, 2013; Macar & Vidal, 2003). As mentioned in the methods section, the ERP analysis was focused on this centroparietal region of interest, between 200 and 800 ms post-offset. The mean amplitude of the six successive 100 ms temporal windows (200-300 ms, 300-400 ms, 400-500 ms, 500-600 ms, 600-700 ms, 700-800 ms) was analyzed, using a two-way repeated-measures ANOVA, with duration (7) and task (2) as within-subject factors. The complete ANOVA tables per temporal window are reported in supplementary material 1.

**Figure 4:**
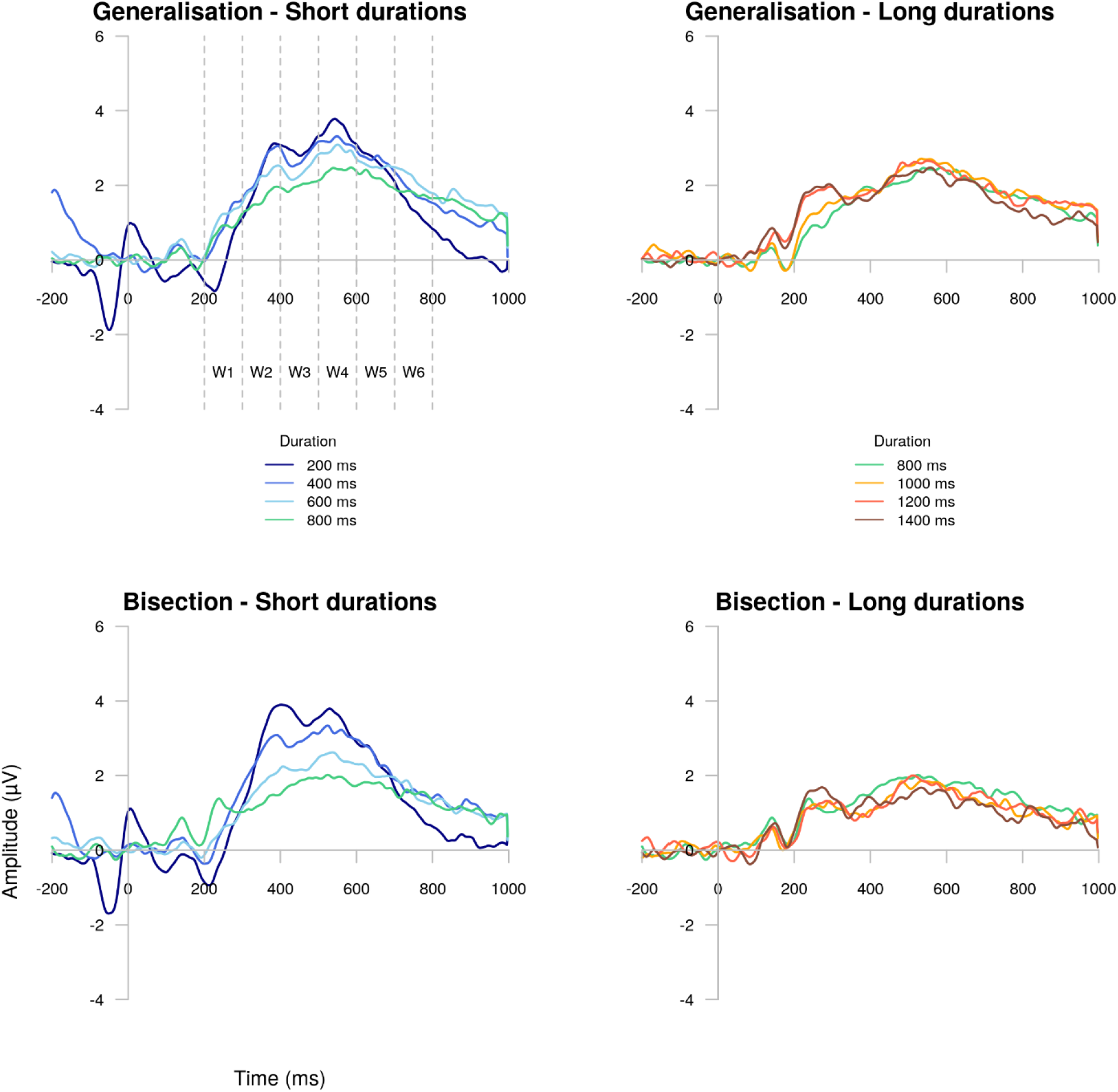
Centroparietal ERPs evoked at the end of each of the seven presented durations. Between 200 ms and 800 ms post-offset, a late positive component (LPC) was apparent. The left panels represent the short durations and the right panels the long ones. The generalisation is shown in the top panels and the bisection in the bottom.

**Figure 5:**
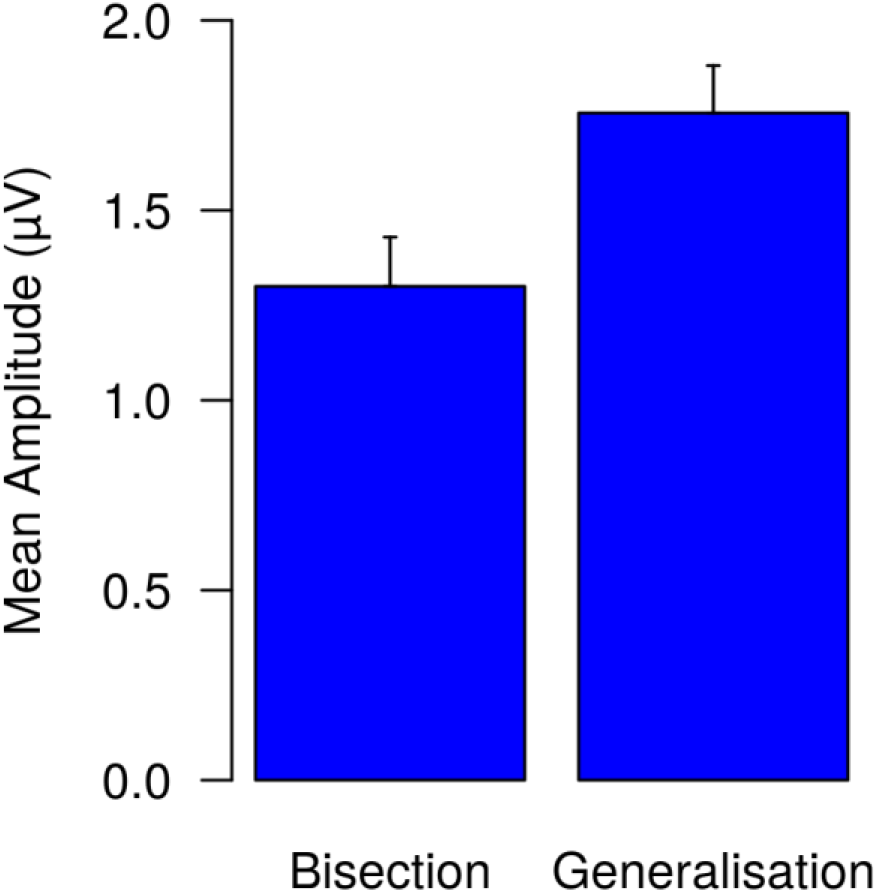
Effect of the task on the LPC mean amplitude between 500 ms and 600 ms after the offset of stimuli.

The effect of the task was significant between 200-300 ms (F(1,17) = 5.37 ; p = 0.033 ; ηp^2^ = 0.24), 300-400 ms (F(1,17) = 5.27 ; p = 0.035 ; ηp^2^ = 0.24), 700-800 ms (F(1,17) = 5.4 ; p = 0.033 ; ηp^2^ = 0.24) and approached significance between 500-600 ms (F(1,17) = 4.26 ; p = 0.05 ; ηp^2^ = 0.20) and 600-700 ms (F(1,17) = 4.16 ; p = 0.06 ; ηp^2^ = 0.19) after stimulus offset. Each time, the mean amplitude of the centroparietal signal was larger in the generalization task than in the bisection one Stimulus duration also affected ERPs between 200-300 ms (F(6,102) = 9.71 ; p < 0.001 ; ηp^2^ = 0.36), 300-400 ms (F(6,102) = 8.37 ; p < 0.001 ; ηp^2^ = 0.33), 500-600 ms (F(6,102) = 11.53 ; p < 0.001 ; ηp^2^ = 0.4) and 600-700 ms (F(6,102) = 3.97 ; p = 0.14 ; ηp^2^ = 0.19). In the first temporal window (200-300 ms), the mean amplitude was higher for longer durations (Figure 6).

**Figure 6:**
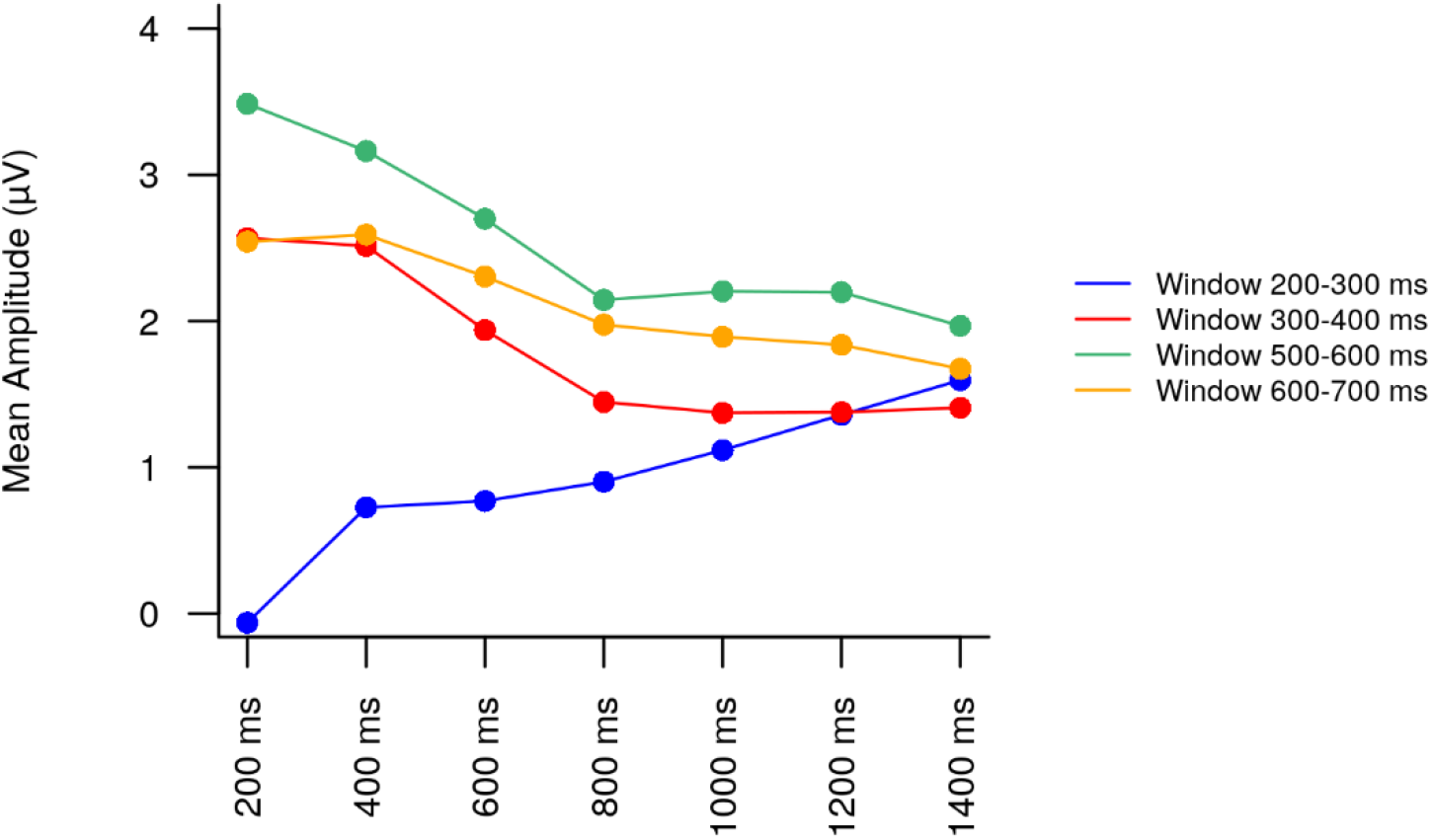
Effect of the presented duration on the LPC mean amplitude in the four temporal windows where the effect was significant : 200-300 ms post-offset (blue), 300-400 ms (red), 500-600 ms (green), 600-700 ms (blue).

Conversely, the pattern was different for the other temporal windows: the amplitude was reduced for longer durations, until it reached a plateau at the 800 ms duration. In the 400-500 ms temporal window, this effect seemed to be modulated by the task used, as revealed by the interaction between duration and task factors (F(6,102) = 2.62 ; p = 0.07 ; ηp^2^ = 0.13). Repeated-measures ANOVA showed that the duration effect was more pronounced in the bisection task (F(6,102) = 9.32 ; p < 0.001 ; ηp^2^ = 0.35) than in the generalization one (F(6,102) = 3.49 ; p = 0,003 ; ηp^2^ = 0.17) in the same window (Figure 7).

**Figure 7:**
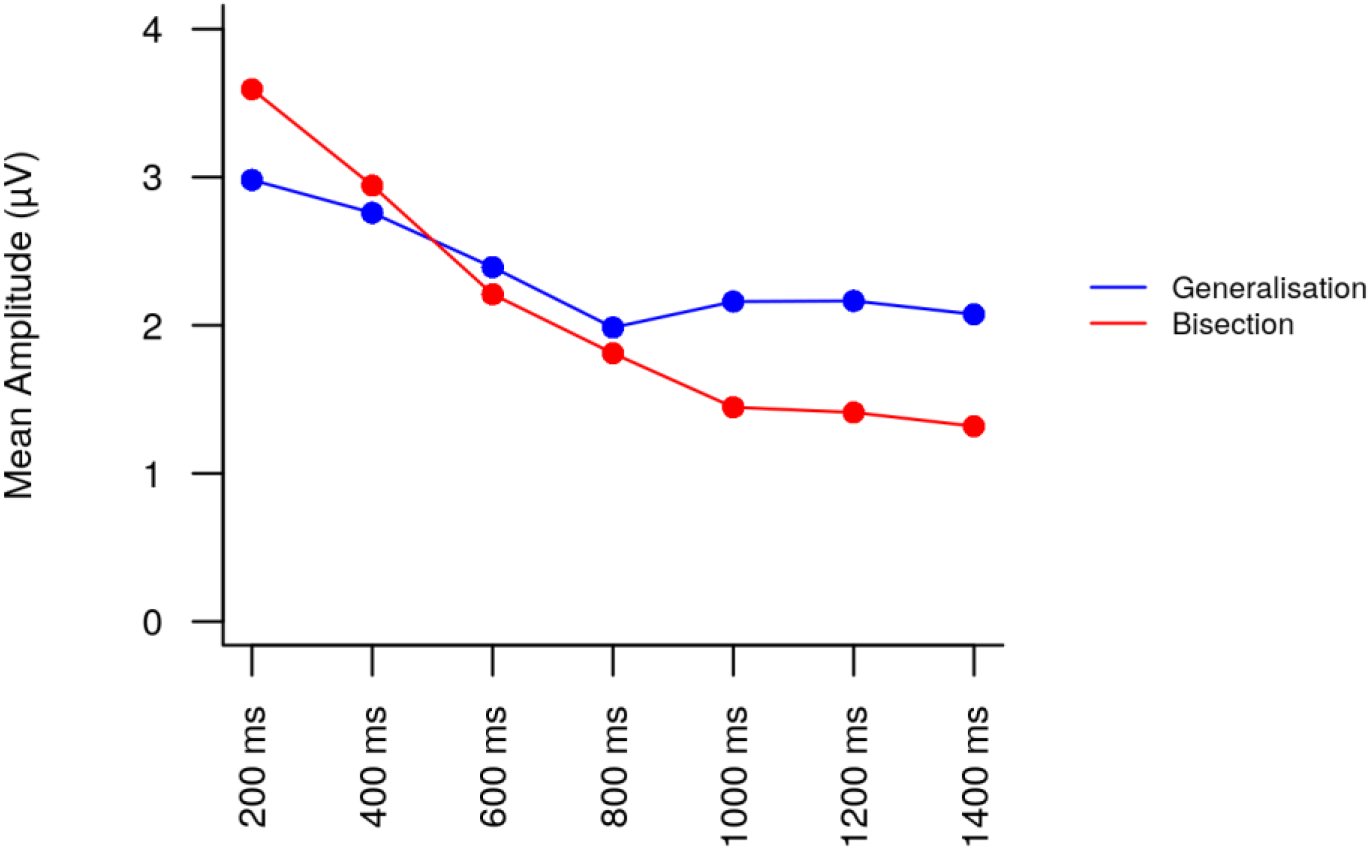
Effect of the presented duration as a function of the paradigm on the LPC mean amplitude, in the 500-600 ms post-offset temporal window.

## 4. Discussion

This study directly investigated differences in decision-making between categorization and identification of durations. We designed a bisection task (categorization) and a generalization one (identification), based on the same temporal stimulus set: 200 to 1400 ms. Two differing hypotheses were tested. On one hand, the bisection task could be more cognitively demanding because it relied on at least two durations in reference memory (the short and long standards). On the other hand, the bisection task could be less cognitively demanding due to the suggestion that there are in fact no standards stored in reference memory in this task (Droit-Volet & Rattat, 2007; Ogden, MacKenzie-Phelan, Mongtomery, Fisk, & Wearden, 2018), while the generalization is based on storage and retrieval of a standard interval. We found several behavioral and ERP results indicating that categorizing durations (bisection) mobilizes fewer resources than identification (generalization). First, participants responded faster at the end of the long durations in the bisection task than in the generalization one. Second, after stimulus offset, centroparietal ERPs revealed that generalization evoked higher amplitudes. This effect interacted with stimulus duration in a temporal window between 400 and 500 ms post-offset. In both tasks, amplitudes decreased as a function of the duration to reach a plateau at the 800 ms presented duration (the median of the intervals distribution), but this effect was much more marked in the bisection task. Third, durations affected also the centroparietal ERPs, independently of the task. Between 200 ms and 300 ms post-offset, ERPs amplitude increased in function of duration. Between 300 ms and 600 ms post-offset, ERPs amplitude decreased in function of duration to a plateau at about 800 ms presented duration.

### 4.1. The behavioral benefits of the bisection task over the generalization one

First of all it should be noted that in the generalization task, the peak of «yes, it is the standard duration » response occurred at 800 ms and 1000 ms, at or near the true standard duration. This demonstrates that the task performance was good. Moreover, the generalization gradient was asymmetric: at a given temporal distance from the standard duration, a longer interval was more confused with the standard than a shorter one. This is in line with what is usually observed in generalization tasks (e.g.Ferrara et al., 1997; Paul et al., 2011). Models of temporal generalization performance suggest that this asymmetry is due to the decision processes that humans use when performing this task (Wearden, 1992). In the bisection task, the bisection point was 722 ms, slightly closer to the geometric mean (676 ms) than to the arithmetic mean (800 ms) of the duration distribution set used and this is concordant with previous studies reporting that the bisection point is located around the geometric mean of durations presented (Allan & Gibbon, 1991), or between the arithmetic and geometric means of the standards or the entire stimulus set..

As the participants were instructed to respond as fast and as accurately as possible, it is quite striking to note that they systematically responded after the end of duration presentation, when in fact they could have responded before the end of long durations as the decision could be made as soon as the standard duration (generalization) or the bisection point (bisection) had elapsed. This unexpected result suggests that participants added an additional task, implying that they were trying to respond as fast as possible at the end of the presented duration, as in a foreperiod paradigm (Los, 2010). Consistent with this hypothesis, RTs decreased as a function of the stimulus duration, revealing that participants used the passage of time to encode the probability of the offset given that it had not yet occurred (Anna C Nobre, Correa, & Coull, 2007). The longer the duration is, the more likely is the offset of the stimulus subtending it is, and the faster the reaction time is. Hence, it seems that at the same time participants completed an explicit timing task (bisection or generalization) and an implicit one, where they used the elapsed duration of the stimulus to improve their reaction time.

Reaction time analysis demonstrated that responses were faster for long durations in the bisection task than in the generalization one. In the case of these durations, participants were able to make a decision before the end of the stimulus. However, as noted above, participants waited the offset of the stimulus to respond as quickly as possible. Therefore, the interaction between task and duration factors reveals that preparation was better for the bisection task for long durations. Decisions could have been made earlier in the bisection than in the generalization for these stimuli, allowing partcipants to better anticipate the stimulus offset. A possible explanation for this advantage for bisection could be in the type of response given by the participants. The bisection task requires responding as to whether the stimulus was more similar to the short or long standards. As duration increases, the probability of responding that the duration is short decreases. Thus, the longer the duration, the more confident the participant is about a « long » response. In contrast, the generalization task required the subjects to judge whether or not the stimulus was the standard. Here, time passing does not facilitate the decision between the two possible responses. However, since this was the case in the bisection paradigm, this could have caused a reduction in decision time in this task.

### 4.2. The bisection task engages fewer resources than the generalization one

Centroparietal ERP data revealed a long positive component (LPC) between 200 ms and 800 ms after stimulus offset. The mean amplitude between 200 ms and 400 ms post-offset on one hand, and between 500 ms and 800 ms post-offset, on the other hand, was higher in the generalization task than in the bisection one. This is in line with previous reports showing that the temporal processing continues after the end of duration presentation (Kononowicz et al., 2016). In particular, similar long positive components were found on centroparietal or frontocentral areas after the end of durations (Giovannelli et al., 2014; Tarantino et al., 2010). These components were linked to attentional and working memory resources allocated to temporal processing. This suggests that the decreased amplitude in the bisection task means that this paradigm engages globally fewer resources than the generalization one, and is concordant with a recent study reporting that bisecting durations could be easier than identifying them (Ogden et al., 2018). According to these authors, this could happen because the bisection does not engage reference memory for short and long standard durations, contrary to what has been proposed by models of bisection (Wearden, 1991, 2004). Instead, a temporal criterion, either based on the mean of all the durations encountered in the task (in the “normal” bisection case) or derived in some way from local duration judgements (in the “episodic” case), would serve in order to bisect intervals (Droit-Volet & Rattat, 2007; Mendoza, Méndez, Pérez, Prado, & Merchant, 2018), implying that this task is less demanding than previously thought.

An interaction occurred between task and duration between 400 ms and 500 ms after stimulus offset on the LPC. In each paradigm, the amplitude decreased with increasing stimulus duration to a plateau reached at about the 800 ms presented duration. However, this phenomenon was much more marked in the bisection task than in the generalization one. In the bisection paradigm, the mean amplitude is higher for short durations and was not very marked for longer durations. This extends the behavioral results and interpretation about RT. Indeed, ERP results in this temporal window indicate that short durations recruit more attentional and working memory resources than long durations. As noted above, the decision seems to be facilitated for long durations due to the passing of time. Indeed, as time passes, the short response is less likely, making easier to choose a long response, implying that short durations have higher cognitive demands than long ones. In the generalization paradigm, the mean amplitude slightly decreases as the stimulus duration increases, but it not as marked as in bisection. In this case, the passing of time cannot help the participant to make a decision, as participants have to decide if the duration is the standard, not whether it is long.

### 4.3. The presented duration moderates the temporal processing after the stimulus extinction

Independently of the paradigm performed by the participants, short durations evoked higher LPC amplitudes than long durations, between 300 ms and 600 ms after the stimulus offset. In contrast, long stimuli evoked higher centroparietal components than the short ones between 200 ms and 300 ms after the stimulus offset. As previously reported, temporal processing after the end of a duration is modulated by the presented duration (Kononowicz & van Rijn, 2014; Lindbergh & Kieffaber, 2013). When a stimulus is shorter than the standard duration, temporal processing is not over at the end of the stimulus presentation. On one hand, decision making had to take place after the stimulus offset, explaining why we observed an important centroparietal activity for these durations between 300 ms and 600 ms after the stimulus extinction. On the other hand, decision-making can occur before the end of the stimuli that are longer than the standard duration, resulting in a temporal processing ending sooner for longer than for short durations.

## 5. Conclusion

The findings of this study suggest that categorizing durations engages fewer resources than identifying them. In the generalization task, participants had to remember a previously learned standard interval in order to determine whether or not the presented duration was or was not identical to it. As previously suggested (Ogden et al., 2018), there may be no such recruitment of long term memory in the bisection task, easing the cognitive demand. Moreover, bisecting durations implies a very different decision than identifying them. This task allows the subjects to rely on the time passing in order to categorize a duration as long when it actually is long. In contrast, the generalization task does not allow this response facilitation as participants do not have to respond if the duration is long, but whether or not it is the standard. Consequently, this study demonstrates the necessity to explore differences in temporal processing between tasks in order to better apprehend the diverse ways we can perceive time.

